# A Data-driven Surrogate Model for Work Computation of a Periodically Forced Half-Sarcomere

**DOI:** 10.1101/2020.03.09.984310

**Authors:** Samuel H. Rudy, C. David Williams, J. Nathan Kutz, Thomas L. Daniel

## Abstract

Muscle force generation follows from molecular scale interactions that drive macroscopic behaviors and macroscopic processes that influence those at the molecular scale. A particuarly challenging issue is that models at the molecular level of organization are often quite difficult to apply to larger spatial scales. This is particularly true of moleuclar models driven by Monte-Carlo simulations. This challenge of multiscale dynamics requires methods to extract reduced order behaviors from detailed high-dimensional simulations. In this work we present a novel deterministic simulation method yielding accurate predictions of force-length behaviors of contracting muscle sarcomeres undergoing periodic length changes (work loops). The model maintains interpretability by tracking macroscopic state variables throughout the simulation while using data-driven representations of dynamics. Parameters of the data-driven dynamics are learned from trajectories from Monte-Carlo simulations of a half-sarcomere. Our method significantly reduces computational cost by tracking the state of the sarcomere in a course grained set of variables while maintaining accurate prediction of macroscopic level observables and time series for course grained variables. This allows for rapid sampling of the model’s output and builds towards the ability to scale to multiple-sarcomere simulations.

**Author Summary:** We develop a data-driven surrogate model for the dynamics of the half-sarcomere. This model achieves the same behavior with respect to force traces as more sophisticated Monte Carlo models at a substantially lower computational cost. The model is built by finding a course grained description of the full state space of the Monte Carlo simulation and learning dynamical models on the course grained space. Data-driven representations of the dynamics in the course grained space are trained using data from the full model. Data-driven models for forcing are also learned, and the result fed back into the dynamics. In doing so, the model seeks to replicate the effects of filament compliance on macro scale dynamics without explicitly tracking micro scale features. We withhold some input parameter regimes and demonstrate accurate reconstruction of course grained state and force traces using the data-driven model and given only knowledge of the initial condition and input. This work allows for faster computation of the forcing behavior of the half-sarcomere, as well as consistent representations of the course grained state variables. It is therefore promising as a step towards multi-sarcomere or even tissue scale models of skeletal muscle.

## Introduction

Since it was first proposed in 1954 by A.F. Huxley *et al.* [1, 2], the sliding filament model for muscle contraction has inspired myriad experimental and theoretical approaches for understanding the molecular and cellular basis of force generation. The focus of the sliding filament model is based on a sarcomere, the fundamental structural unit of muscle cells. Sarcomeres are subcellular structures arranged in chains along the length of a muscle cell. Each sarcomere, in turn, is composed of overlapping protein filaments running between sequential z-disks. Actin containing thin filaments attach to the z-disks. The thin filaments overlap with thicker myosin containing filaments in the center of each sarcomere. Myosin filaments are, in turn, equipped with crossbridges that act together as ensembles of molecular motors to pull the actin filaments towards the center of the sarcomere, thus shortening its length and driving a sliding motion between the thick and thin filaments (see Figure 1).

**Fig 1.**
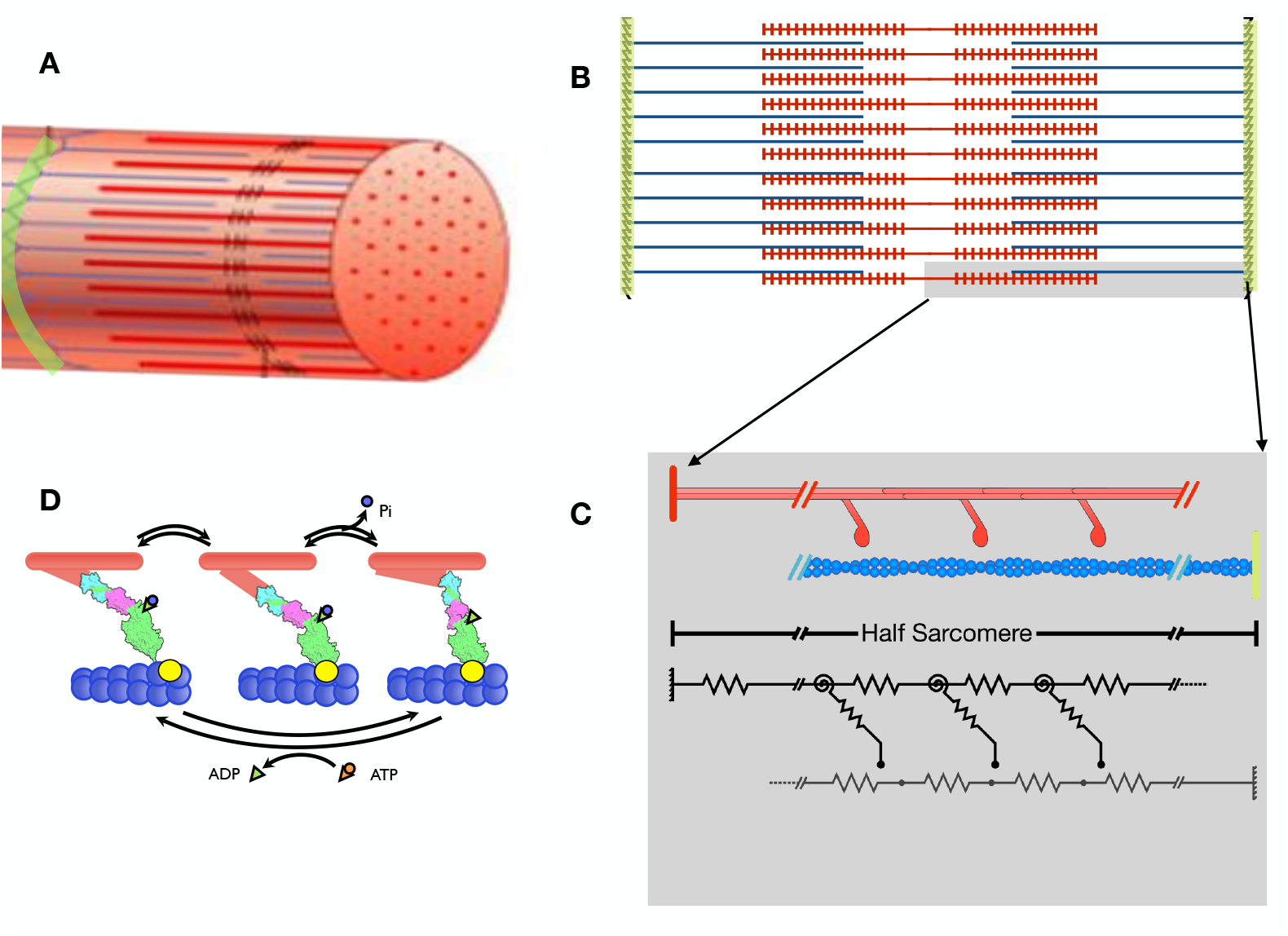
Geometry. Schematic figure for the geometric basis of the models developed by [13, 14]. Muscle cells (A) consist of axially repeating sarcomeres (B) which of which contain thin filaments comprised of double helical arrangements of actin monomers (blue) anchored to z-disks (green). Thin filaments interdigitate with thick filaments (red) that contain myosin motor molecules anchored to the backbone of thick filaments. The model developed in prior studies [8, 9, 13, 14] is based on a three state system with transitions shown in D.

**Fig 2.**
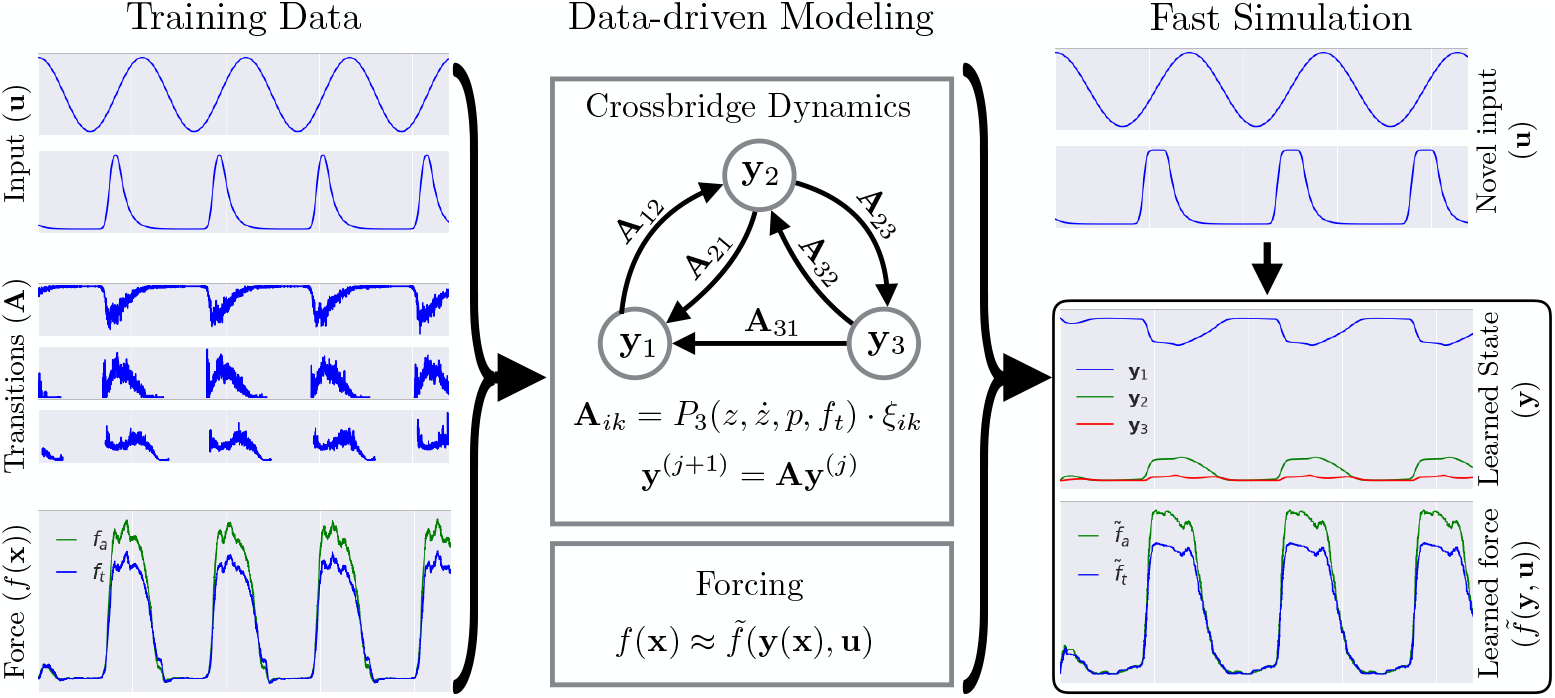
Schematic for method. Schematic figure for data-driven modeling of muscle. (left) data is collected from MC simulation of full system. (middle) transition probabilities are modeled as a Markov matrix having edge weights dependent on external forcing. Measurements (force and tension) are modeled as functions of external input and number of cross bridges in each state. (right) A new instance of external forcing is considered, Markov transition probabilities give states of cross bridges, and force and tension are then computed.

Mathematical models for muscle contraction based on sliding filament theory were first proposed in 1957 [3] and drew largely on ordinary differential equation models that assumed cross-bridges detach and attach to binding sites on the thin filaments according to a first-order kinetic scheme. These and many other early models sought to explain the transduction of chemical energy into mechanical work. With the initial assumptions that actin and myosin filaments were inextensible, the force generated by the sarcomere was computed by simply summing the forces generated by each crossbridge as a function of its local distortion and effective spring constant. Assuming a continuum limit the crossbridge’s density along the length of the myosin filaments and binding sites along actin filaments, one can derive differential equation models for the sarcomere [3].

Following early modeling work that assumed filaments were inextensible, more recent experimental evidence suggested that filaments do in fact exhibit compliance [4, 5]. This discovery had several implications for any mathematical model of the half sarcomere. First, the assumption that each crossbridge behaves independently would not apply, since the distortion of the filament lattice in response to local force generation by one crossbridge would influence the binding probability of other crossbridges.. Additionally, the net axial force would no longer be a simple sum of all forces created by crossbridges since lattice distortion influences the total force. Thus, the local deformation of filaments may lead to coupling between crossbridges via realignment of binding sites due to filament compliance. Indeed, experimental evidence suggests a highly nonlinear relationship between filament overlap, which corresponds to the number of bound crossbridges, and force. [6].

Accounting for filament compliance with partial differential equation (PDE) models is nontrivial, though the original Huxley model has been adapted to include extensible filaments [7]. The adapted PDE based theory, however, does not consider mechanical coupling between crossbridges. Monte Carlo techniques have subsequently been proposed that explicitly track the location of each crossbridge and binding site, as well as crossbridge states [8, 9]. These models started with single filament pair interactions [8] and were extended to consider radial geometry of the sarcomere [9]. In contrast to PDE based models, Monte Carlo simulations have revealed that compliance based mechanical coupling does play a role in crossbridge dynamics. Additionally, similar Monte Carlo models for force generation by cardiac muscle cells have shown how coupling between active sites on thin filamentscan drives emergent dynamics [9, 10]

Macroscopic and tissue scale models of muscle will require large scale coupling of individual sarcomere models. Coupling of small numbers of sarcomeres has been explored in the differential equations model case [10, 11]. Coupled models explicitly including effects from filament compliance require scaling already expensive computational models and are thus infeasible. Recent work has looked at coupled ordinary and partial differential equation models that mimic results of Monte Carlo simulations [12] but has not been extended to the multiple sarcomere case. The prohibitive cost of scaling Monte Carlo based simulations motivates surrogate models that allow for faster computation of the input output dynamics of the Monte Carlo simulations with lower computational expense.

This work constructs a data-driven surrogate model for the half-sarcomere meant to replicate the behavior of the spatially explicit Monte Carlo techniques discussed in [9, 13, 14]. Each simulation of the model is parameterized by an externally imposed length. Binding sites on each actin filament have permissiveness modulated by calcium, which binds to troponin molecules on the filaments inducing a conformal change and allowing binding. In contrast to past methods which represent dynamics as differential equations, we use a discrete model to track the probability of crossbridges being in one of three bound states throughout the simulations. Transition probabilities between states are modeled in a data-driven manner from simulation data using the Monte Carlo technique outlined in [9, 13, 14]. We are specifically interested in the case of periodically lengthened and activated half-sarcomeres. Here we seek to develop accurate reduced order models of force generation by the half sarcomere in response to periodic length changes and a variety of input parameter regimes. These reduced order models, in turn, can be used to bridge dynamics at microscopic scales to mechanical behaviors at macroscopic scales.

## Materials and Methods

In this section we more formally discuss the problem of building a surrogate model for force calculations and provide detailed explanations for the computational techniques used. The section is organized as follows: in the subsection *Mathematical Formulation* we motivate the problem of a reduced cost surrogate model using an abstract notation of the full scale mechanistic model presented in [8] and also discussed in [13, 14]. In the subsection *Data-driven half-sarcomere model* we provide a detailed description of the computational techniques used in this work as well as considerations for model validation. In all that follows we will be discussing a dynamical system evolving in time with discretized increments *t_j_*. The notation •^(*j*)^ will always refer to the quantity • evaluated at time *t_j_*.

### Mathematical formulation

We first consider an abstract motivation of the surrogate model. The half-sarcomere consists of a bundle of actin and myosin filaments on which we impose periodic length changes and calcium based modulation of binding site permisiveness. We denote the state of the half-sarcomere at time *t*, including locations of each cross-bridge, their bound state, and locations of each binding site as 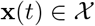. Input parameters, including actin permissiveness and externally induced length changes are denoted by 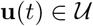. A single step of the mechanistic simulation described in [8] is given by,

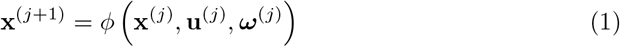

where ****ω****^(*j*)^ ∈ Ω is a random variable capturing the simulation’s stochasticity. In practice, ****ω**** is a number of identical independently distributed uniform random variables used to compute bound states of each cross-bridge. In this work we restrict our attention to periodic input. Let 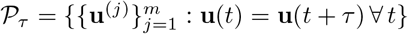 be the space of length *m* discretely sampled time series of *τ* periodic signals and 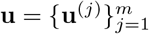. The full length mechanistic simulation in [8] is then a function 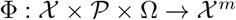 given by,

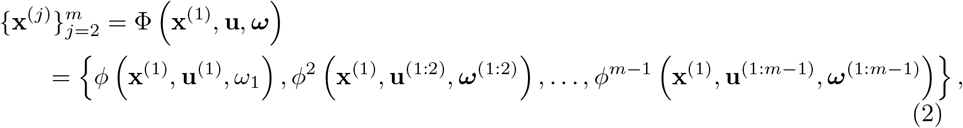

where 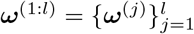 and

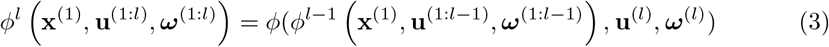

for *l* > 1. Equation (2) allows one to sample trajectories of **x** given a prescribed input parameterization **u**. In this research we are interested in the work generated by the half sarcomere. Letting 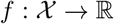 denote the state-dependent force generated by the half-sarcomere, work over one loop of the periodic forcing is,

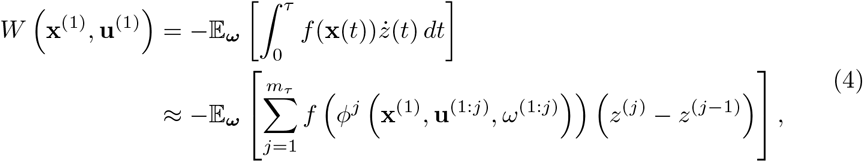

where *z*(*t*) is sarcomere length, *m_τ_* = *τ/dt*, and the negative sign has been added since force is computed in the negative *z* direction. Evaluation for the full model ϕ requires sampling from Ω as well as a complex numerical optimization problem for each step. In some cases, for example when evaluating the work generated by a new parameter regime, it is useful to use a lower fidelity model with decreased computational cost. Critically, any such model must also maintain sufficient resolution to accurately compute the force exerted by the half-sarcomere at each timestep in the simulation. In this work, we find a reduced set of variabes 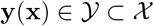, function 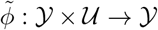, and 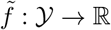 such that,

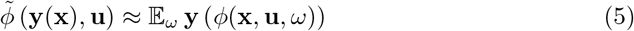

and 

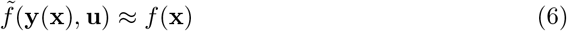

both hold with small residuals for all **x**, **u** in a large training set. That is, we want a dynamical model 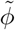 of the course grained space that is consistent with the full model *φ* and a consistent means 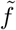 for computing force from the course grained space.

The fact that **u** is in the domain of 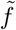 and not *f* implicitly allows for unresolved state variables to affect the force computed from the course grained space, so long as they react quickly to **u**. A prototypcial example is crossbridge length, which is modulated by change in sarcomere length encoded in **u**.

### State space of the mechanistic simulation

We provide a brief description of the state space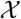 of the mechanistic simulation φ used a high fidelity model and refer the reader to [9, 13, 14] for more detail. The simulation exploits the symmetry of a sarcomere to focus modeling on the span between a single z-disk and m-line, the midpoint of the sarcomere. A grid of actin and myosin filaments is arranged according to the multifilaments geometry discussed in [9]. Spatial location of each crossbridge and binding site are explicitly tracked throughout the simulation. We enumerate the crossbridges by which myosin filament they lie on and their location on that filament, and likewise for binding sites on actin filaments. The state space of the mechanistic simulation is uniquely defined by

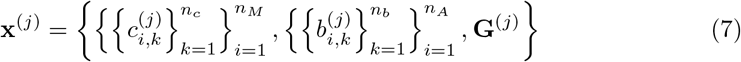

where *n_M_* and *n_A_* are the number of myosin and actin filaments, *n_c_* is the number of crossbridges per myosin filament, and *n_b_* is the number of binding sites per actin filament. Variables *c* and *b* are positions of crossbridges and binding sites, respectively. **G** is an adjacency matrix encoding connections between crossbridges and binding sites. It is defined by

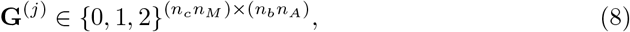

where 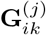 is 1, 2, or 3 if the cross bridge indexed by *i* is unbound, loosely bound, or tightly bound to the binding site indexed by *k* at time *t_j_*. This representation of connectivity is a large overparameterization. Since no more than one crossbridge may attach to a given binding site and a crossbridge may only bind to one site, at most one element of any particular row or column of **G** may be nonzero. Many entries will necessarily be zero due to the simulation’s geometry. However, it will be useful for describing changes in the sarcomere’s state that we aim to capture in the data-driven model.

All simulations in this work are parameterized by three input time series; the length of the half sarcomere, lattice spacing, and actin permissiveness. The last variable models binding site affinity modulated by calcium. These are abbreviated as *z*^(*j*)^, *l*^(*j*)^, and *p*^(*j*)^, respectively and are described by time series,

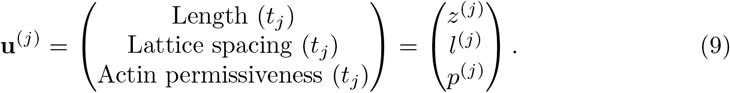

We assume constant volume so that 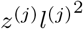 is constant throughout any simulation. Three output variables are recorded throughout the simulation: axial force, radial force, and radial tension. We denote these by

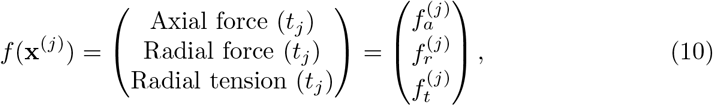

where it is implicitly understood that each is dependent on **x**^(*j*)^. Radial force exists in the plan perpendicular to the sarcomere and is generally very close to zero. We are primarily concerned with accurate computations of axial force throughout a simulation.

The training data is composed of a number of runs of the mechanistic model using many parameter regions. We let 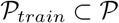denote the set of parameter regimes used in the training data. Since we do not vary the initial condition **x**^(1)^, each run is characterized by its input parameters **u** and random variable ****ω**** ∈ Ω^*m*^. We denote the collection of random variables used for each parameter regime **u** as Ω_**u**_⊂ Ω^*m*^.

### Data-driven half-sarcomere model

We first consider the problem of finding a good set of course grained coordinates 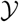 on which to construct a data-driven surrogate model. In many fields of computational physics such as fluid dynamics, there are established means of finding reduced order bases on which to construct reduced models [15]. The problem here is less straight forward, since many variables tracked in the mechanistic simulation are categorical and we cannot generally assume that dynamics are constrained to a low dimensional space. We therefore rely on domain knowledge of the half-sarcomere to determine 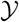.

The fundamental force generating mechanisms in the actin-myosin interaction are the cross bridges, which in the mechanistic Monte-Carlo model exist in one of the states; free, loosely bound, or tightly bound. In our simulations, the mechanistic model accounts for 720 crossbridges. Our surrogate model averages over the number of crossbridges in each state to achieve a significant reduction in dimensionality. We will show that simply tracking the fraction of cross-bridges in each state, along with input parameters and forcing, is sufficient to satisfy equations (5) and (6). The course grained state variables **y** and approximation of force *f* used in this work are,

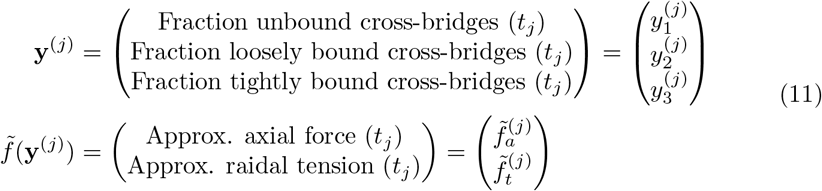

Radial force is omitted from the data-driven model since it is expected to fluctuate in a small region about zero and is not relevant for computing work. Radial tension was found to be important for computing the dynamics of the cross-bridge states.

#### Dynamics model in course grained space

The internal state of the half-sarcomere is represented only with the fraction of crossbridges in each of the three bounds states. It is therefore natural to construct a dynamics model based on transition rates between these states. A canonical model for state and input dependent transition rates is given by,

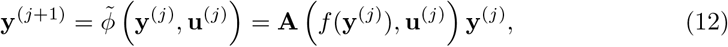

where the constraint 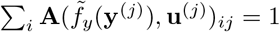 enforces that the total number of crossbridges in maintained. Since crossbridges do not communicate with each other, it seems intuitive that the dynamics should be linear in **y**. Indeed, the mechanistic model *f* computes transition probabilities for individual crossbridges based on distance to binding site and binding site permissiveness. However, **u** alone seems to be insufficient to accurately compute transition probabilities and we include radial tension. Individual transition probabilities are modeled using polynomial regression on features including radial tension and input. Specifically,

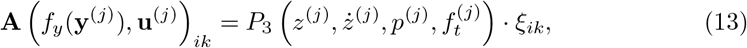

where *P_q_* indicates the set of polynomial features of order less than or equal to *q* in the features and *ξ_ik_* is a learned weight vector. The term **A**(*f_y_*(**y**^(*j*)^), **u**^(*j*)^)_*ik*_ is the probability of a crossbridge transitioning from state *i* to state *k* during the timespan (*t_j_, t_j_*_+1_].

We impose several constraints on the matrix **A**. Transitions from unbound (state 1) to tightly bound (state 3) are prohibited in the mechanistic model so **A**_13_ is defined to be zero. We also need to ensure each column sums to one and that entries are non-negative. Empirical evidence from the mechanistic simulation suggests that the diagonal entries of **A** corresponding to the probability that a crossbridge does not change state are much larger than off-diagonal. We train only off-diagonal entries and subtract those from each column sum to obtain the probability of no transition. To ensure non-negativity we project each transition probability onto [0, 1] after applying (13). This strategy could in theory still result in a negative probability if the two learned off-diagonal entries sum to greater than one, but this has not been observed and could easily be flagged during a simulation.

#### Training the dynamics model

Weight vectors *ξ_ik_* are trained using data from the mechanistic model. This requires computing transition probabilities between states at each timestep of each of the simulations used as training data. It is be helpful to consider a simpler notion of crossbridge connectivity than **G** where we ignore binding sites and focus only on the bound state of each crossbridge. The time dependent collection crossbridge state vector 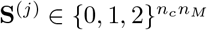 is given by,

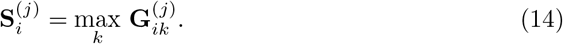

This relates nicely to the course grained variables through the relation,

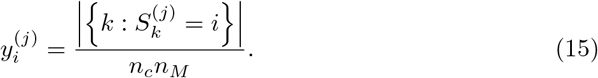

Empirical transition probabilities for each step of the simulation are given by, 

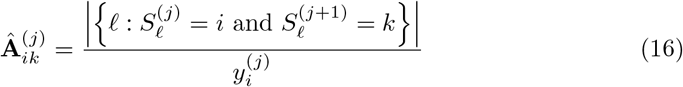

where 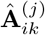 is left undefined when *y_i_* = 0. The matrix of empirical transition probabilities 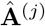 will vary across simulations depending on initial condition **x**^(1)^, input **u**, and random variable ****ω****.

Each of the *ξ_ik_* is trained using the collection of all timesteps in each simulation across all parameter regimes considered in the training dataset. The numerator of (16) is a draw from the binomial distribution 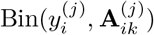 and thus has mean 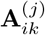 and variance 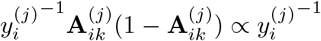. A maximum likelihood estimate of *ξ*_*ik*_ should weigh each observation where 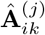 by the approximate precision 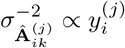. However, this method does not yield predictive results. The denominator of (16) is highly correlated with the actin permissiveness *p*^(*j*)^ and other input parameters. Indeed, the mechanistic simulation generally has perioidic intervals in which *y*_2_ = *y*_3_ = 0. Instead, we threshold observations where 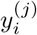 is below a fixed tolerance *y_min_* = 5 and weight the remaining observations equally.

Since the cubic polynomial kernel may result in a more ill-conditioned linear system we impose an *L*^1^ penalty to encourage parsimonious weight vectors, as is common in many physical applications [16–18]. To maintain a tractable size for the linear system even with many trials of each parameter region, we average over the empirical transition rates and radial tension within each parameter regime. The resulting cost function is given by,

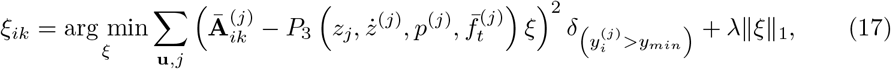

where 

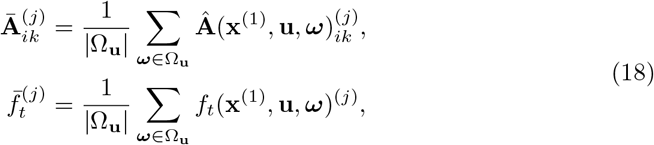

and *δ* is a Kronecker delta function indicating when there is sufficient sample size to trust the observation. Each monomial term in the range of *P*_3_ is normalized across the training dataset so that regularization is independent of measurement units. Optimization for (17) is performed in the Scikit Learn python library [19] using 10-fold cross validation to find regularization term *λ*.

### Forcing model in course grained space

Force in the mechanistic simulation is computed by summing over forces generated by individual crossbridges. However, there are properties of the crossbridges not encoded in the averaged bound states. Each crossbridge acts as a spring which is stretched according to the length between it’s base on a myosin filament and binding site. This distance changes according to input parameters. Therefore, the data-driven model for forcing is constructed as a function of the averaged cross-bridge states as well as the input parameters **u**.

Specifically we construct 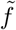 as an ensemble of regression trees using gradient boosting [20] as implemented in Scikit Learn [19]. The gradient boosting algorithm sequentially constructs an emsemble of regression trees [21]. The resulting 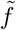 is of the form,

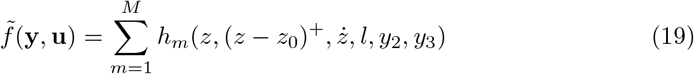

where each *h_m_*(**y**, **u**) is a regression tree and *M* = 100. The term (*z* − *z*_0_)^+^ is the positive component of the length minus it’s temporal mean and allows for distinct functional forms between when length is greater or less than its average. Actin permissiveness and the number of free crossbridges have been omitted since they do not have direct effect on forcing. We use the squared error loss function so that each tree *h_i_* is trained on the residual error of the sum over all previous trees. Forcing and state fractions used in training 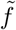 are averaged over runs having the same input parameters. The optimization for each tree is given by,

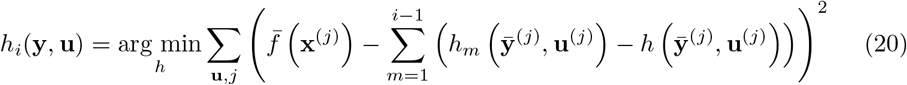

where 

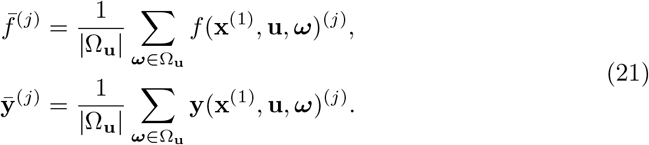

Each regression tree *h_m_* is a piecewise constant function that splits the domain along variables that result in minimal variance of the target function in each half of the resulting split domain. This is performed recursively until a maximum depth of 5 is achieved or variance reduction in the splitting is below a threshold.

There are several advantages to this approach. The learned function 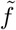 is composed of a large ensemble of weak learners which allows the model to be accurate without suffering from overfitting. The resulting 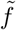 yields more accurate predictions than other commonly used supervised learning techniques, including regularized polynomial regression and neural networks. Regression trees also constrain predictions to lie within the same range as the training data so novel inputs to 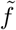 will not result in extreme values.

### Simulating new data

The data driven model allows us to compute forcing given new input parameters 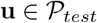. Trajectories from the data driven simulation are represented by,

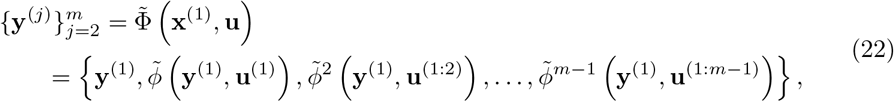

where *Y*^(1)^=*Y*(*x*^(1)^) The work done by the half-sarcomere over one period of oscillation is computed using the data driven approximation of force as, 

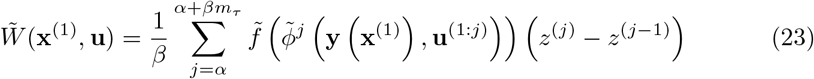

where *β* = 3 is an integer number of periods to average over and *α* = *m* − *βm*_*τ*_ is a burn in time to allow for decay of any initial transient onto a periodic orbit.

## Results

The dataset used to train the data-driven model consists of 20 sample trajectories from each of 150 unique parameter regimes in 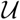. It is split into 135 training parameter regimes and 15 for model evaluation. Input parameters are altered by changing the period of length changes [45. 45, 55. 55, 71.43 ms], phase offset of calcium activation as fraction of length cycle period [0, 0.1, 0.2,…, 0.9], and half life of calcium activation [12, 15, 18, 21, 25 ms]. In this section we compare results of the Monte Carlo simulation to those from the data-driven simulation on the parameter regimes reserved for testing. All results from the data-driven model are computed only from the initial condition of the mechanistic simulation using the same inputs **u**. Therefore, accuracy is shown not only for point wise computations of transition probabilities, but of the full length time series.

Figure 3 shows the results for a single parameter regime of the eight learned transition probabilities, 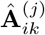, and empirical transition probabilities averaged over all 20 trials, 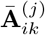. Empirical transition probabilities are only shown in the plot where they are defined for each of the sample trajectories, with gaps where the denominator of Eq. 16 is less than *y_min_*. Substantially increased variance in certain regions is likely due to low populations in the pre-transition state during those periods.

**Fig 3.**
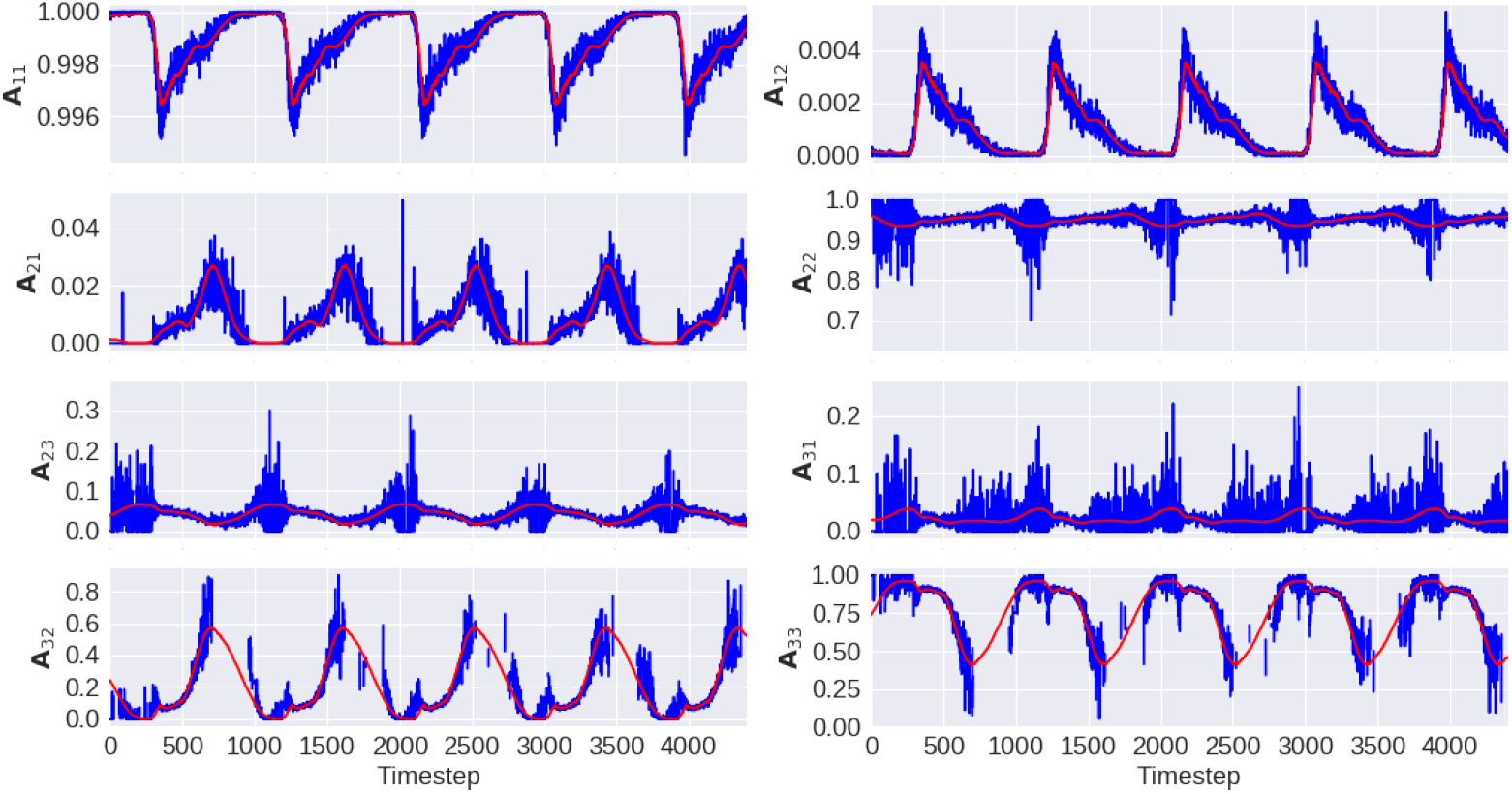
Transition probabilities. Example of empirical transition probabilities from mechanistic model (averaged over 20 trials) in blue and computed transition probabilities from data-driven model in red for input regime in testing data. Missing values for mechanistic data indicate undefined transition probability due to empty state.

The crossbridge state densities resulting from the transition probabilities in Fig. 3 are shown in Fig. 4. Blue curves and regions indicate the mean and one standard deviation band of observed cross-bridge state densities across all trials of the Monte Carlo simulation for this parameter regime. While slight deviation is observed in the data-driven model, it largely mimics the behavior of the Monte Carlo simulation.

**Fig 4.**
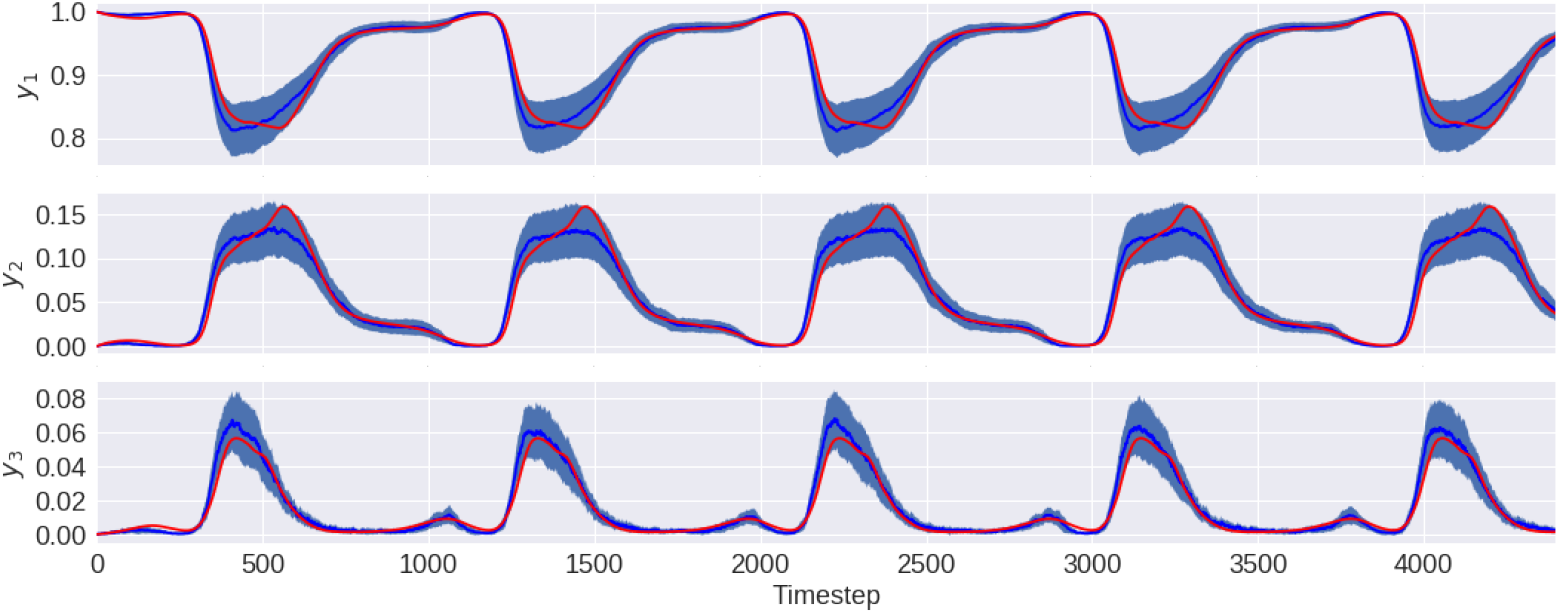
Crossbridge States. Example of crossbridge state averages for the mechanistic model in blue and data driven model in red for parameter regime in testing data. Mechanistic data is averaged over 20 runs with shaded region indicating one standard deviation.

Finally, for the same parameter regime shown in figures 3 and 4, Fig. 5 shows the axial force and radial tension computed using the data-driven model, 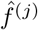, in red, and from the mechanistic simulation, 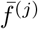, in blue. Shaded regions indicate one standard deviation bands.

**Fig 5.**
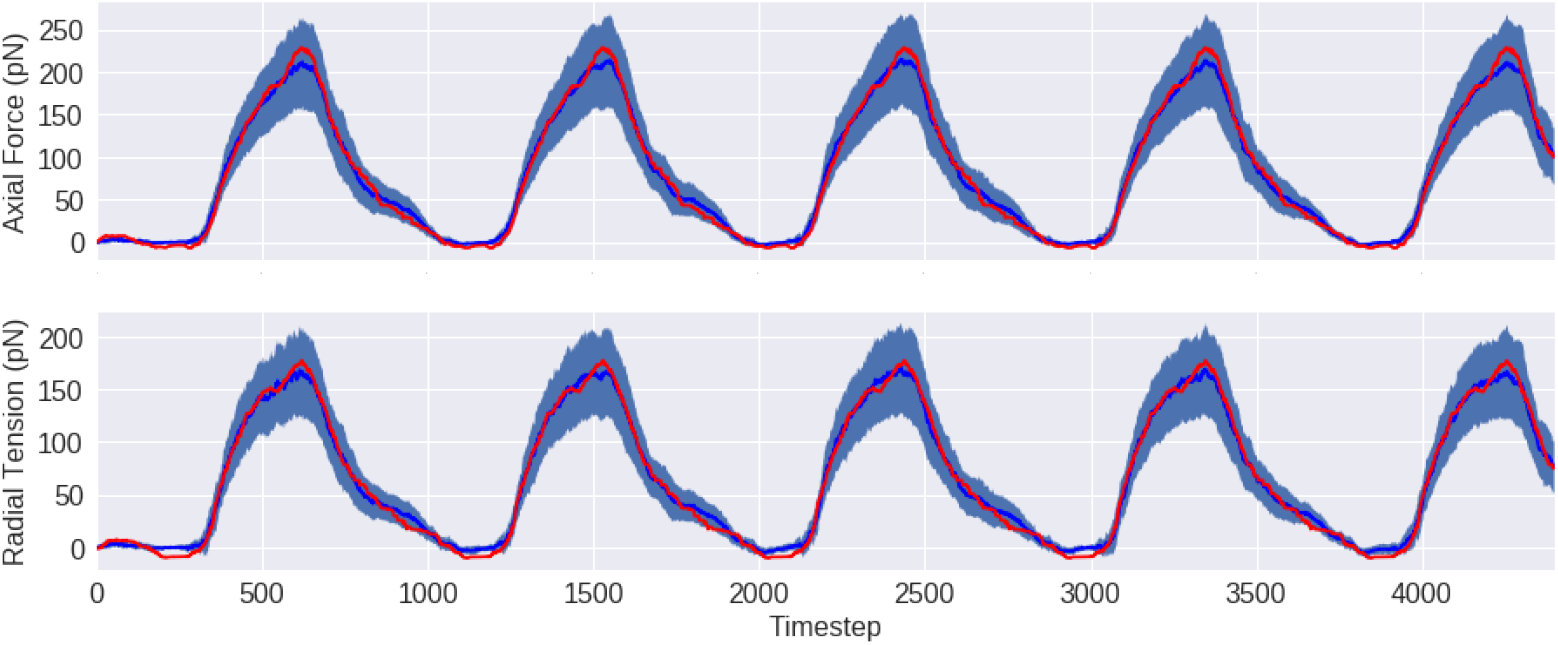
Forcing. Example of forcing for the mechanistic model in blue and data driven model in red for parameter regime in testing data. Mechanistic data is averaged over 20 runs with shaded region indicating one standard deviation.

Work loops for each of the 15 testing regimes are shown in Fig. 6. Red trajectories are generated using the data-driven methods and blue trajectories are given by the average force from the Monte Carlo simulation, 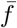. A cursory inspection reveals that the data-driven model fails to capture high forcing in the case where there are multiple loops and significant forcing occurs at minimal length. These cases seem to be correlated with high calcium activation during the initial stage of sarcomere shortening.

**Fig 6.**
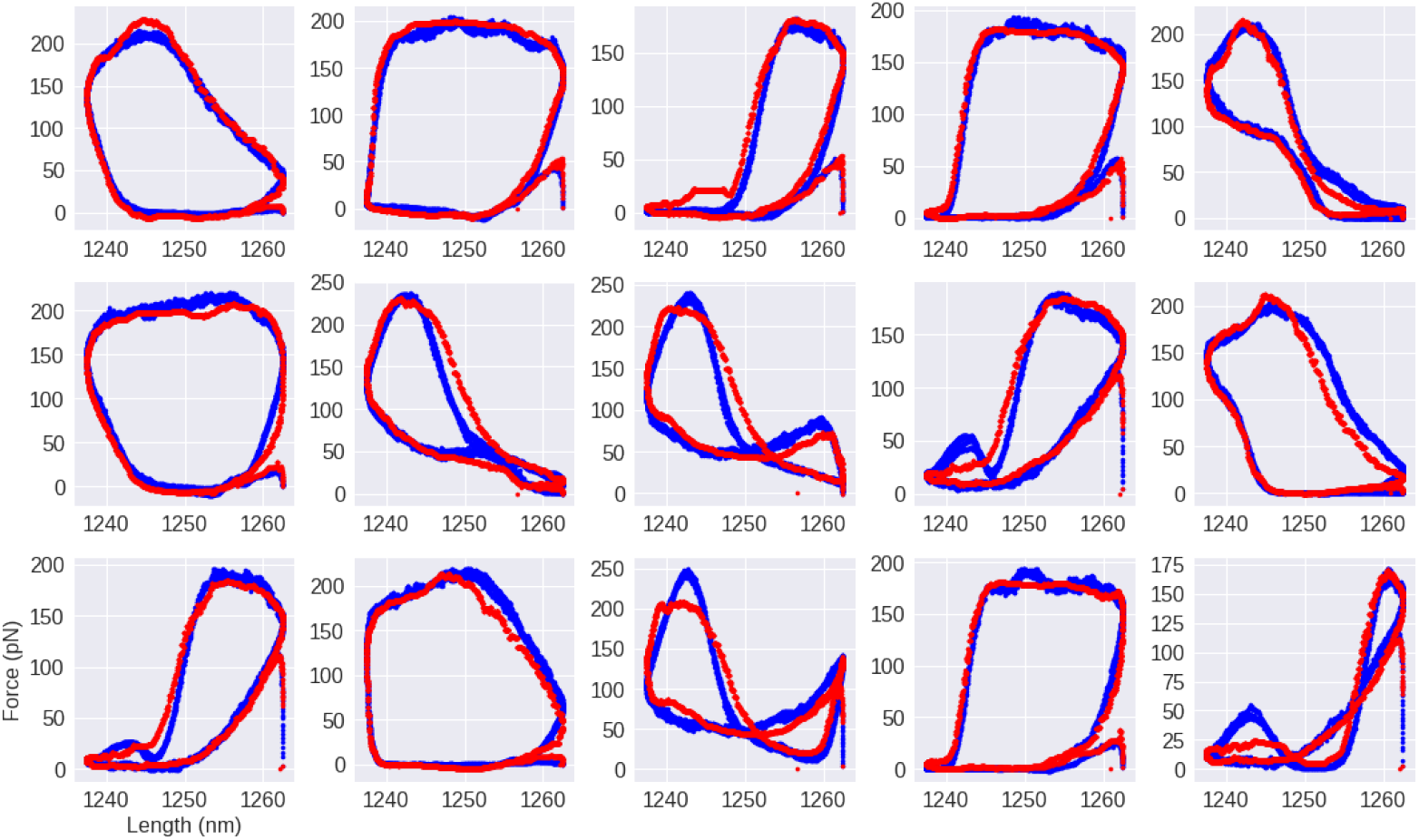
Workloops for testing data. Computed workloops for testing data according to mechanistic model in blue and data driven model in red. Each plot corresponds to an input regime not seen during the training procedure.

Figure 7 summarizes the key results in this chapter regarding work loop computations. We compare work from the mechanistic (Monte Carlo) model with that from the data-driven model. Parameter regimes included in the training data are shown in blue, and those from the testing data in red. We see that work approximated by the data-driven model for both training and testing regimes closely adheres to what is observed in the Monte Carlo model.

**Fig 7.**
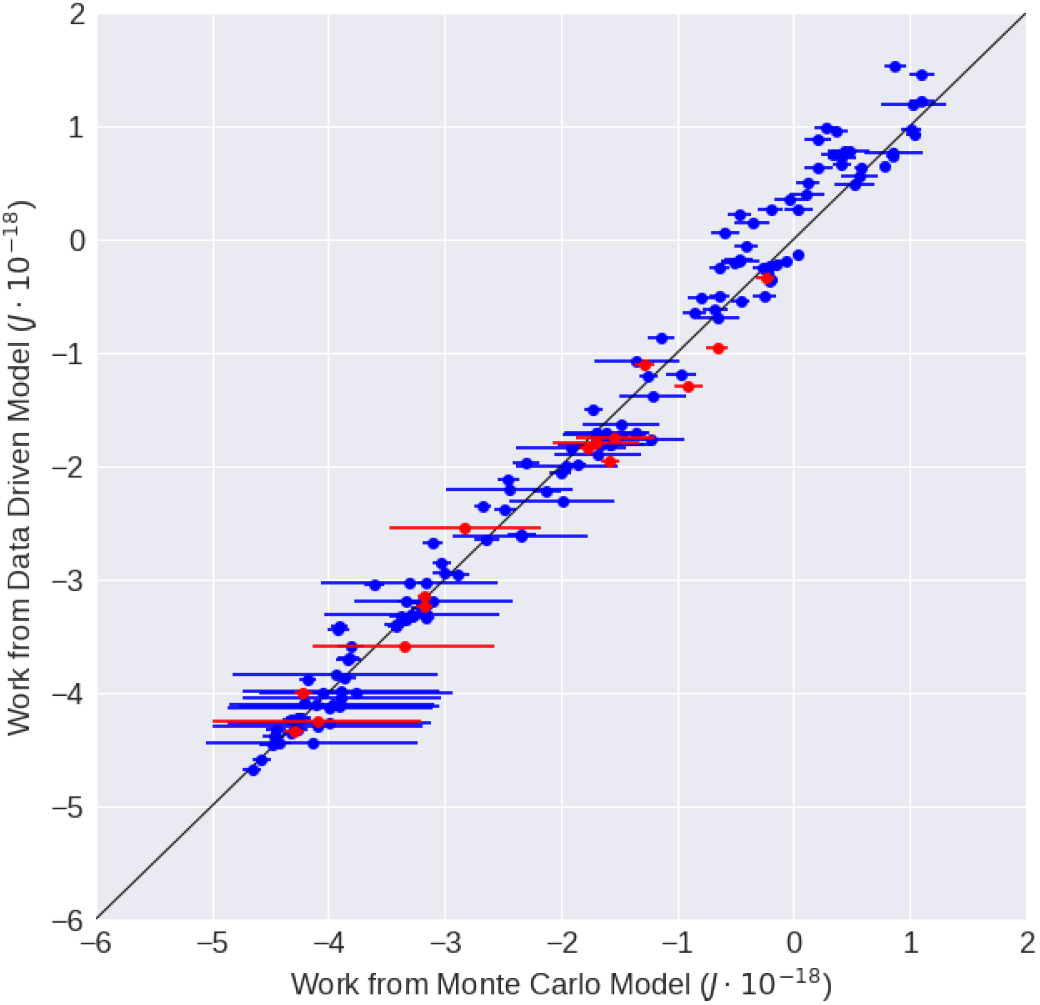
Work. Computed work in mechanistic model on x-axis and data driven model on y-axis. Blue data points correspond to input in training data while red is testing. Error bars indicate one standard deviation around mean value observed over 20 trials of the mechanistic model.

## Discussion

We have presented a novel data-driven technique for modeling the half-sarcomere under periodic length changes and activation. The data-driven model is trained on data from and mimics the behavior of the Monte Carlo techniques developed in [8, 9] at a significantly decreased computational expense. In particular, force traces and work loops are largely consistent when the two models are compared.

The current work offers several paths for future research. While methods presented here replicate dynamics on the course grained variables and force traces given new input parameters, several regimes in the testing set indicate that some features are missed. Increases in axial force during the contracting phase of the sarcomere are underestimated in some cases, even though crossbridge state densities are accurate.

This indicates that the data-driven model for force could be improved through different parameterizations of 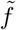 or by including finer variables in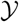. However, such work could come at the expense of increased model complexity and could significantly increase the challenge in finding dynamics 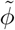 in the course grained space.

One goal of the current work to build towards a scalable model that may be used to run simulations of many sarcomeres coupled end to end. In this case, periodic length changes imposed on the string of sarcomeres may not imply periodicity on each sarcomere, so the model would need to be tested on a larger space on inputs. Furthermore, physiologically accurate models at a more macroscopic scale would require considering this force as acting on a set of sarcomeres with masses moving in a viscous medium. Tuning for drag coefficients may be a highly nontrivial problem.

We have constructed the framework in this work to demonstrate that it is feasible to replicate the behavior of the Monte Carlo simulation with a fast deterministic system, but do not claim that the method presented is optimal. It is common for reduced order models of Markovian dynamical systems in physics to lose the Markov property and thus require a memory term for accurate representation of dynamics in the reduced space [22]. There is reason to believe that the dynamics in course grained variables developed in this work would benefit from incorporating non-Markovian terms, though data-driven simulation would then require initialization with a short time seriesrather than only requiring the initial condition.

To facilitate future work and in the interest of reproducible research, all data and code used for this paper has been made publicly available under an MIT license on GitHub at https://github.com/snagcliffs/data driven sarc.

## Acknowledgments

This work was supported by grants to TLD from the Army Research Office W911NF-14-1-0396, the National Institutes of Health P30AR074990, the Washington Research Foundation and the Joan and Richard Komen University Endowment.

